# Diversity of *Mycobacterium tuberculosis* across evolutionary scales

**DOI:** 10.1101/014217

**Authors:** Mary B O’Neill, Tatum D Mortimer, Caitlin S Pepperell

## Abstract

Tuberculosis (TB) is a global public health emergency. Increasingly drug resistant strains of *Mycobacterium tuberculosis* (*M.tb*) continue to emerge and spread, highlighting adaptability of this pathogen. Most studies of *M.tb* evolution have relied on ‘between-host’ samples, in which each person with TB is represented by a single *M.tb* isolate. However, individuals with TB commonly harbor populations of *M.tb* numbering in the billions. Here, we use analyses of *M.tb* genomic data from within and between hosts to gain insight into influences shaping genetic diversity of this pathogen. We find that the amount of *M.tb* genetic diversity harbored by individuals with TB can vary dramatically, likely as a function of disease severity. Surprisingly, we did not find an appreciable impact of TB treatment on *M.tb* diversity. In examining genomic data from *M.tb* samples within and between hosts with TB, we find that genes involved in the regulation, synthesis, and transportation of immunomodulatory cell envelope lipids appear repeatedly in the extremes of various statistical measures of diversity. Many of these genes have been identified as possible targets of selection in other studies employing different methods and data sets. Taken together, these observations suggest that *M.tb* cell envelope lipids are targets of selection within hosts. Many of these lipids are specific to pathogenic mycobacteria and, in some cases, human-pathogenic mycobacteria. We speculate that rapid adaptation of cell envelope lipids is facilitated by functional redundancy, flexibility in their metabolism, and their roles mediating interactions with the host.

## Author Summary

Tuberculosis (TB) is a grave threat to global public health and is the second leading cause of death due to infectious disease. The causative agent, *Mycobacterium tuberculosis* (*M.tb*), has emerged in increasingly drug resistant forms that hamper our efforts to control TB. We need a better understanding of *M.tb* adaptation to guide development of more effective TB treatment and control strategies. The goal of this study was to gain insight into *M.tb* evolution within individual patients with TB. We found that TB patients harbor a diverse population of *M.tb*. We further found evidence to suggest that the bacterial population evolves measurably in response to selection pressures imposed by the environment within hosts. Changes were particularly notable in *M.tb* genes involved in the regulation, synthesis and transportation of lipids and glycolipids of the bacterial cell envelope. These findings have important implications for drug and vaccine development, and provide insight into TB host pathogen interactions.

## Symbols and Abbreviations

Multidrug-resistant (MDR), Extensively drug-resistant (XDR), Nucleotide diversity (π), Watterson’s theta (ƟW), Tajima’s D (TD), ratio of counts of non-synonymous variants per non-synonymous site to synonymous variants per synonymous site (*πN/πS*), Between-host Tajima’s D (BHTD), Within-host Tajima’s D (WHTD), Polyketide Synthase (PKS), Mycobacterial membrane protein Large (MmpL), phthiocerol dimycocerosates (PDIM), sulfolipids (SL), polyacyl trehalose (PAT), diacyl trehalose (DAT), phenolic glycolipid (PGL), novel complex polar lipids synthesized by Pks6 (POL), and mannosyl-β-1-phosphomycoketides (MPM)

## Introduction

*Mycobacterium tuberculosis* (*M.tb*) causes over nine million new cases of tuberculosis (TB) per year and is estimated to infect one-third of the world’s population [1]. The emergence of increasingly drug resistant strains of *M.tb* [2] demonstrates the bacterium’s ability to adapt to antibiotic pressures, despite limited genetic diversity [3]. Prior research has identified the influence of bottlenecks, population sub-division, and purifying selection on genetic diversity of *M.tb* circulating among human hosts [4–7]. In these studies, each TB patient was represented by a single *M.tb* strain isolated in pure culture. However, individuals with TB harbor a large population of *M.tb* cells for a period of months to years, which raises the possibility of significant diversification of bacterial populations over the course of individual infections.

There are few studies of within-host evolution of *M.tb*. One example is a study of the transposable element IS6110 marker that found multiple lines of evidence suggestive of positive selection on *M.tb* populations within hosts [8]. Advances in sequencing technologies have since enabled detailed, genome-wide studies of the evolution of intra-host populations of both pathogenic and commensal microbes [9–22]. Whole-genome sequencing (WGS) studies of natural populations of *M.tb* have focused primarily on the emergence of drug resistance [23–26]. Here, we use analyses of genetic data from within and between patients with TB to characterize *M.tb* variation across evolutionary scales. We find that overall diversity of *M.tb* populations within hosts can vary dramatically and identify candidate genetic loci for *M.tb* adaptation during infection.

## Results and Discussion

### Genome-wide variation

We quantified genetic diversity of five within-host populations of *M.tb* (Table 1) using data from three published studies. The data we selected from the original studies were directly comparable with respect to sequencing platform and other factors we observed to affect diversity (*see Methods* and S1 Table). The data set includes samples susceptible to all first line drugs (A0, E0) as well as INH-monoresistant, multidrug-resistant (MDR) and extensively drug-resistant (XDR) samples (Table 1). HIV status was not reported in any of the primary studies. *M.tb* lineage 2 (East Asian, Patients A, B, C and D) and lineage 4 (Euro-American, Patient E) are included in this sample; patient origin was reported as China (Patients A, B, and C), Abkhazia (Patient D) and “Eastern Europe” (Patient E). We used PoPoolation software [27] to estimate two standard measures of genetic diversity [nucleotide diversity (π) and Watterson’s theta (ƟW); see Inset Box for descriptions] from the pooled within-host *M.tb* genomic data. It is challenging to distinguish rare genetic variants from sequencing errors in pooled genomic data: PoPoolation implements methods that account for the effects of sequencing errors on low frequency variant allele calls [27,28]. In order to eliminate effects of coverage differences on diversity, we sub-sampled the genomic data to a uniform 50X coverage (see *Methods*).

**Table 1.**
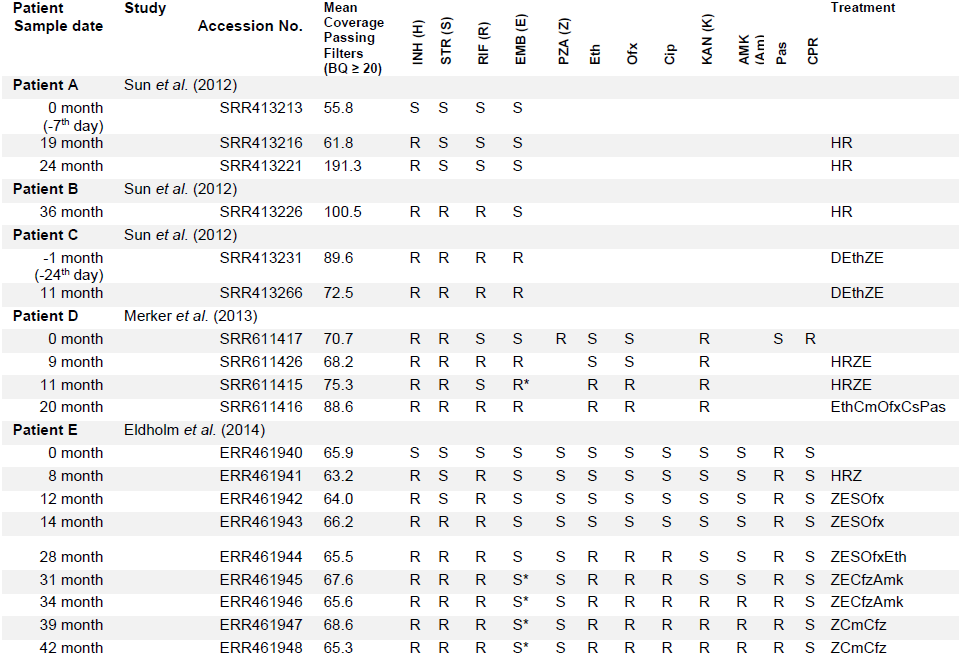
Within-host samples of *Mycobacterium tuberculosis*. Sample dates and resistance profiles are from the respective publications; sample timing is in reference to treatment initiation. Treatment at sampling is listed. Resistance profile abbreviations: R=resistant; S=susceptible; blank=not reported; *indicates more details presented in the original paper. Drug abbreviations: INH (H), isoniazid; STR (S), streptomycin; RIF (R), rifampicin; EMB (E), ethambutol; PZA (Z), pyrazinamide; Eth, ethionamide; Ofx, ofloxacin; Cip, ciprofloxacin; KAN (K), kanamycin; AMK (Am), amikacin; Pas, para-aminosalicylic acid; CPR (Cm), capreomycin; D, Dipasic (isoniazid aminosalicylate); Cs, Cycloserine; Cfz, clofazimine.

| Patient<br>Sample date | Study | Accession No. | Mean<br>Coverage<br>Passing<br>Filters<br>(BQ ≥ 20) | INH (H) | STR (S) | RIF (R) | EMB (E) | PZA (Z) | Eth | Ofx | Cip | KAN (K) | AMK (Am) | Pas | CPR | Treatment |
| --- | --- | --- | --- | --- | --- | --- | --- | --- | --- | --- | --- | --- | --- | --- | --- | --- |
| **Patient A** | Sun *et al.* (2012) |  |  |  |  |  |  |  |  |  |  |  |  |  |  |  |
| 0 month<br>(-7<sup>th</sup> day) |  | SRR413213 | 55.8 | S | S | S | S |  |  |  |  |  |  |  |  |  |
| 19 month |  | SRR413216 | 61.8 | R | S | S | S |  |  |  |  |  |  |  |  | HR |
| 24 month |  | SRR413221 | 191.3 | R | S | S | S |  |  |  |  |  |  |  |  | HR |
| **Patient B** | Sun *et al.* (2012) |  |  |  |  |  |  |  |  |  |  |  |  |  |  |  |
| 36 month |  | SRR413226 | 100.5 | R | R | R | S |  |  |  |  |  |  |  |  | HR |
| **Patient C** | Sun *et al.* (2012) |  |  |  |  |  |  |  |  |  |  |  |  |  |  |  |
| -1 month<br>(-24<sup>th</sup> day) |  | SRR413231 | 89.6 | R | R | R | R |  |  |  |  |  |  |  |  | DEthZE |
| 11 month |  | SRR413266 | 72.5 | R | R | R | R |  |  |  |  |  |  |  |  | DEthZE |
| **Patient D** | Merker *et al.* (2013) |  |  |  |  |  |  |  |  |  |  |  |  |  |  |  |
| 0 month |  | SRR611417 | 70.7 | R | R | S | S | R | S | S |  | R |  | S | R |  |
| 9 month |  | SRR611426 | 68.2 | R | R | R | R |  | S | S |  | R |  |  |  | HRZE |
| 11 month |  | SRR611415 | 75.3 | R | R | S | R* |  | R | R |  | R |  |  |  | HRZE |
| 20 month |  | SRR611416 | 88.6 | R | R | R | R |  | R | R |  | R |  |  |  | EthCmOfxCsPas |
| **Patient E** | Eldholm *et al.* (2014) |  |  |  |  |  |  |  |  |  |  |  |  |  |  |  |
| 0 month |  | ERR461940 | 65.9 | S | S | S | S | S | S | S | S | S | S | R | S |  |
| 8 month |  | ERR461941 | 63.2 | R | S | R | S | S | S | S | S | S | S | R | S | HRZ |
| 12 month |  | ERR461942 | 64.0 | R | S | R | S | S | S | S | S | S | S | R | S | ZESOfx |
| 14 month |  | ERR461943 | 66.2 | R | R | R | S | S | S | S | S | S | S | R | S | ZESOfx |

|  |  |  |  |  |  |  |  |  |  |  |  |  |  |  |  |
| --- | --- | --- | --- | --- | --- | --- | --- | --- | --- | --- | --- | --- | --- | --- | --- |
| 28 month | ERR461944 | 65.5 | R | R | R | S | S | R | R | R | S | S | R | S | ZESOfxEth |
| 31 month | ERR461945 | 67.6 | R | R | R | S* | S | R | R | R | S | S | R | S | ZECfzAmk |
| 34 month | ERR461946 | 65.6 | R | R | R | S* | S | R | R | R | R | R | R | S | ZECfzAmk |
| 39 month | ERR461947 | 68.6 | R | R | R | S* | S | R | R | R | R | R | R | S | ZCmCfz |
| 42 month | ERR461948 | 65.3 | R | R | R | S* | S | R | R | R | R | R | R | S | ZCmCfz |

##### Description of statistical measures used in the text.

We used two measures to quantify genetic diversity in the samples included in our study. Nucleotide diversity (π) is the proportion of loci at which sequences differ in pairwise comparisons (π/site is reported in the text). Watterson’s theta (ƟW) is an estimate of the population mutation rate (the product of the effective population size and mutation rate). It is based on the number of segregating sites in a sample of sequences. Segregating sites are loci at which differences are found in one or more sequences. We report ƟW/site in the text.

Tajima’s D (TD) computes the difference between scaled average numbers of pairwise differences and segregating sites. In a neutrally evolving population of constant size, TD is expected to be zero. Many neutral and selective influences can cause TD to deviate from zero. For example, selection against deleterious mutations (‘purifying’ selection), population expansion, and past selective sweeps (in which an advantageous mutation rapidly increases in frequency) can cause TD to decrease. TD can increase as a result of population sub-division and selection that maintains diversity in the population.

*πN/πS* is the ratio of counts of non-synonymous variants per non-synonymous site to synonymous variants per synonymous site. Stably maintained amino acid polymorphisms (*πN/πS* > 1) may indicate diversifying selection or local sweeps (selection for advantageous mutations under a regime of restricted migration).

We compared the pooled *M.tb* within-host data to a globally diverse sample of 201 ‘between-host’ genomes from the seven major lineages, where each individual with TB is represented by a single isolate of *M.tb* (S2 Table). These data were originally reported in Comas et al [29]. Drug resistance phenotypes were not reported in the original study. We examined the between-host alignment at 1004 loci with a reported association with drug resistance [30] and found that 37 of these loci had segregating polymorphisms. We conclude that the between-host dataset is likely to include both susceptible and drug resistant isolates.

Estimates of *M.tb* diversity within hosts were very sensitive to coverage and platform; measures of diversity from some samples were also sensitive to base quality score and minor allele count threshold (S3 File, S4 Table). Patients D and E were similar in that both developed progressive drug resistance during treatment, and therapy was tailored to results of extended drug susceptibility testing (DST). Patient E’s treatment outcome was eventually successful; treatment outcome of Patient D was not reported. For a given set of parameters, estimates of diversity from Patients D (*M.tb* lineage 2) and E (*M.tb* lineage 4) were very similar to each other, and stable across serial samples (Fig. 1 and S4 Table). Estimates from Patient E were slightly more sensitive to the minor allele count threshold, suggesting this patient harbored more rare variants than Patient D.

**Fig. 1.**
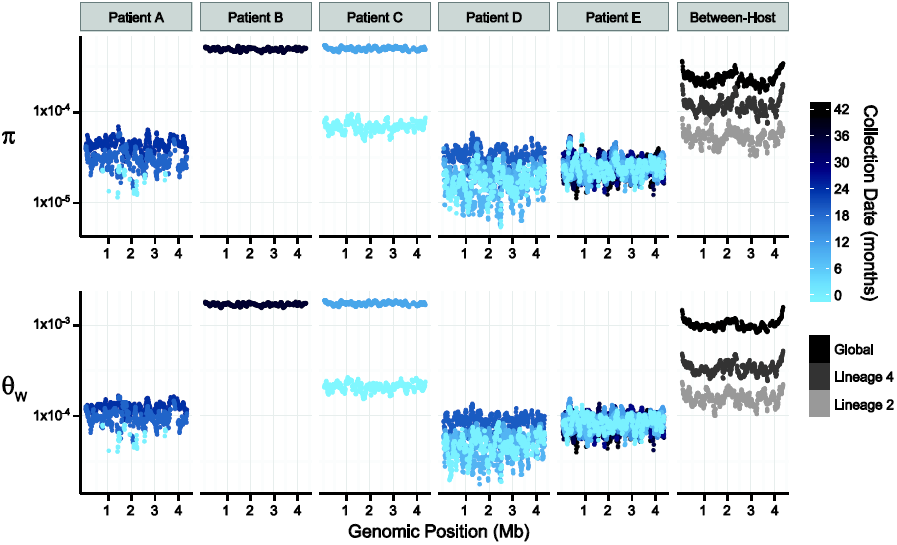
Patterns of nucleotide diversity (π) and Watterson's theta (ƟW) across the *Mycobacterium tuberculosis* (*M.tb*) chromosome. Sliding-window analyses were performed using 100-Kb windows with a step-size of 10-Kb on uniformly subsampled alignments of each sample (50X sequence coverage); plotted values are the mean of each window across 9 replicate subsamplings. Chromosomal coordinates reflect the genomic positions of the reference strain H37Rv, against which pooled-sequence reads were mapped. Temporal samples of each patients’ *M.tb* population are colored to reflect the sample collection date as shown in the legend; global and lineage-specific estimates are colored as indicated in the legend.

Patient A (*M.tb* lineage 2) was described as non-adherent during standard therapy of drug-susceptible disease; the second and third samples from this patient exhibited INH-monoresistance. Patient A was treated with two months of four-drug treatment, followed by 27 months of INH and RIF (i.e. only one drug to which the bacteria were susceptible). Measures of diversity from the *M.tb* population of Patient A were similar overall to those from Patients D and E, but showed an increasing trend over the sampling period (Fig. 1). Increasing diversity could indicate loss of control of the infection. Patient A defaulted treatment, and no further samples were reported after the sample collected at 24 months into treatment.

Patients B and C had MDR TB. Extended DST results were not reported for these patients, and both died. *M.tb* diversity of pre-terminal samples from patients B and C was extremely high relative to the other intra-host samples (Fig. 1). This difference in diversity could be due to a technical issue, such as relatively high sequencing error rates in these data. In this case, we would expect that application of more stringent quality filters would decrease observed diversity. Application of a higher base quality threshold did not result in loss of relative diversity of these samples. Differences between samples from Patients B, C and the others were, however, less marked when the minor allele count threshold was increased (S4 Table). This suggests that high diversity of B36 and C11 was driven by rare variants. Excess rare variants are a hallmark of an expanding population; diversity of pre-terminal samples from Patients B and C could reflect expansion of the *M.tb* population in the final phases of an uncontrolled infection. Another possibility is that terminal progression of their TB infections involved extensive breakdown of lung tissue allowing sampling of previously inaccessible *M.tb* sub-populations.

TB patients included in our within-host data set were culture positive for prolonged periods: this is atypical for settings with well-functioning TB control programs and these patients are unlikely to be broadly representative. Further studies of within-host diversity of TB patients from a range of settings and with a variety of clinical presentations are needed to fully characterize *M.tb* adaptation within hosts. Nonetheless, these data delineate some interesting patterns of within-host diversity. It is striking that *M.tb* within-host diversity is so similar across patients from three independent studies (Patients A, D, and E). *In vitro* estimates of *M.tb*’s mutation rate vary substantially according to lineage [31], yet we did not observe an obvious effect of lineage on diversity in comparisons of *M.tb* populations from lineage 2 (Patients A and D) and lineage 4 (Patient E). This could reflect differences between *in vitro* and *in vivo* mutation rates, or perhaps other parameters are more important in shaping overall patterns of diversity within hosts. Diversity of *M.tb* lineage 2 was lower than lineage 4 at the between-host scale (Fig. 1), whereas lineage 2 isolates have been observed to evolve more quickly in the lab.

Our data set includes four within-host samples that were collected prior to the initiation of treatment (A0, C1, D0, and E0). We did not observe a substantial decrease in *M.tb* diversity following initiation of treatment in any of these patients (A19, C11, D9, and E8). It could be that TB treatment does not affect the average number of pairwise differences and segregating sites of resident *M.tb* populations. Alternatively, measures of *M.tb* diversity may change in response to treatment in TB patients whose sputum cultures convert more quickly than the patients in our study. Our study included two *M.tb* samples with susceptibility to all first line agents (A0 and E0). Diversity of these samples was not markedly different from samples with a drug resistant phenotype.

In summary, our results suggest that *M.tb* lineage, initiation of TB treatment and drug resistance do not have strong impacts on diversity measures for within-host *M.tb* populations. Disease severity, on the other hand, appears to have marked effects on *M.tb* diversity.

### Genes and gene categories with distinct patterns of variation across evolutionary scales

Several studies have identified distinct patterns of variation at *M.tb* genetic loci associated with different functions [4,32]. Such patterns of variation may reflect distinct regimes of natural selection, heterogeneity of mutation rates, or other influences. We sought to identify bacterial genetic loci with extreme patterns of variation at within- and between-host scales; these are candidate loci for *M.tb* adaptation during transmission and infection.

For each gene with data that passed our quality thresholds (see *Methods*), we quantified π and ƟW using methods that account for sequencing error [27]. We used the same approach and estimated two statistics designed to capture deviations from neutral patterns of variation: Tajima’s D (TD, [33]) and the ratio of non-synonymous changes per non-synonymous site (*πN*) to synonymous changes per synonymous site (*πS*) (see Inset Box). We disregarded genes lacking either synonymous or non-synonymous variation in comparisons of *πN/πS*. Since neutral forces such as population growth can affect patterns of variation, we compared relative values of statistics in each within-host sample, where all genes are likely to have the same demographic history.

Natural selection can lead to population differentiation when the relative fitness of genotypes varies among environments; empirical outlier analyses for loci with extreme measures of population differentiation are commonly used to identify candidates of positive selection [34–36]. Treating each serial sputum sample from an individual patient as a distinct *M.tb* population, we calculated pairwise FST values for polymorphic sites covered by at least 10 reads with a minimum minor allele count of 6 (pooled across all samples from a patient) with PoPoolation2 [37]. In order to reduce biases introduced by variable coverage, we conditioned our analysis on the ability to detect a significant change in allele frequency between samplings using a two-sided Fisher’s exact test as previously proposed [38] (*see Methods*).

### Patterns of variation among genes involved in lipid metabolism

In order to identify candidate groups of genes under selection, we examined the extreme tails (≤ 5^th^ percentile, ≥ 95^th^ percentile) of π, Ɵw, and TD for enrichment of specific functional categories. We did not observe any consistent patterns of enrichment in genes with extreme values of π, Ɵw, or the upper tail of TD (S5 Table). There was, however, a striking pattern of enrichment among genes with extremely low values of TD across evolutionary scales (Fig. 2). The Tuberculist [39] categories “lipid metabolism” (LIP), “conserved hypotheticals” (CHP), and the Clusters of Orthologous Groups (COG) [40] category “secondary metabolite biosynthesis, transport, and catabolism” (COG:Q) are significantly enriched in the low tail of the distribution of gene-wise TD values of most samples, and all patients, in the within-host studies, as well as in the global between-host sample (Fig. 2; S5 Table).

**Fig. 2.**
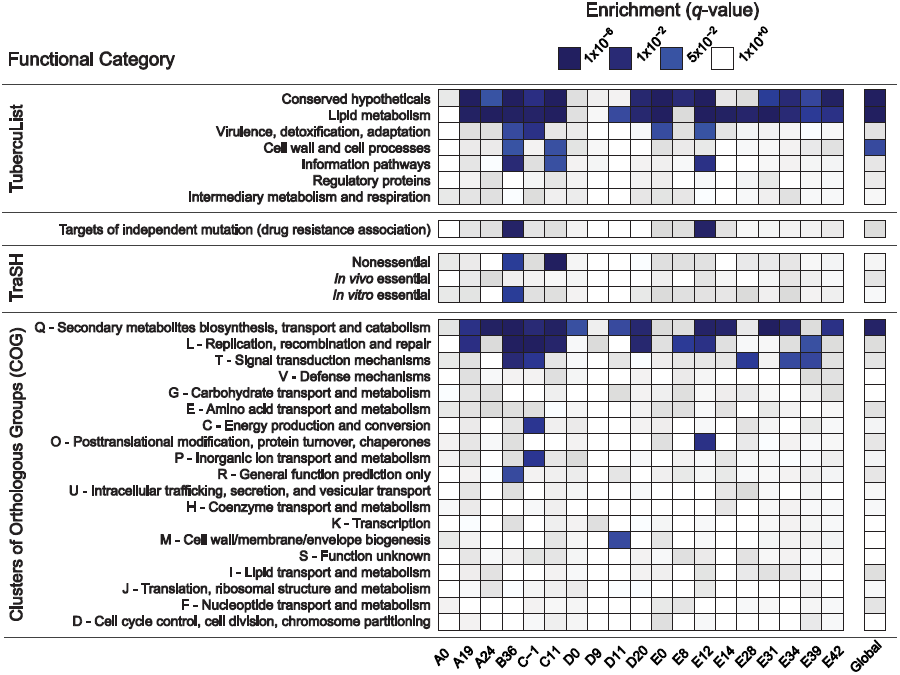
Enrichment of annotation categories among genes with extreme negative values of Tajima’s D (TD). Within each sample (labeled at the bottom of the heatmap), gene-wise TD values were compared and the bottom 5% of genes in the distribution were tested for overrepresentation of functional categories using a two-sided Fisher’s exact test. To account for multiple hypothesis testing, a false discovery rate of 5% was used and the resulting *q*-values are displayed on a continuous scale with varying shades of blue indicating significance at the 0.05 level. The manually curated TubercuList “conserved hypotheticals” and “lipid metabolism” categories [39], as well as the computationally predicted COG:Q “secondary metabolites biosynthesis, transport and catabolism” [40] are notable for their consistent enrichment at both the within- and between-host scales.

To clarify whether the same genes are driving the enrichment of LIP, COG:Q, and CHP categories across evolutionary scales, we examined the overlap of genes in the bottom 5% tail of the between-host distribution with the bottom 5% tails of the TD distributions from within-host samples: all 181 genes in the bottom tail of the between-host TD distribution are also in the tail of at least one within-host sample’s TD distribution.

Low values of within- and between-host TD in COG:Q (234 genes in category) are driven in part by genes that are also categorized as LIP (272 genes in category). Eighty-eight genes overlap between the two classification schemes, of which 42 are in the bottom tail of at least one within-host sample, and 12 are in the bottom tail of the between-host, gene-wise TD distribution. Intriguingly, in a study that analyzed *M.tb* RNA-Seq data over the course of a TB infection, COG:Q was also found to be enriched among genes whose expression was down-regulated [25]. Eldholm et al did not look for differences in expression of TubercuList categories, so we do not know whether LIP and/or CHP were differentially expressed in their study.

Our results indicate that influences producing distinct patterns of variation among genes in LIP, CHP and COG:Q categories are found across a range of human genetic backgrounds and environments. The fact that expression of COG:Q genes appear responsive to the environment within hosts provides further evidence in support of the hypothesis that they are targets of adaptation during infection.

Systematic errors in sequencing and mapping at LIP, CHP and COG:Q loci could potentially also produce unusual patterns of genetic variation. In order to investigate this possibility, we compared base and mapping quality scores of variants in COG:Q, CHP and LIP with genome-wide values. We did not identify significant differences in quality scores among genes in these categories, suggesting that extreme patterns of variation are not driven by errors in sequencing and processing of sequencing data (S6 Figure).

Mutation rate variation is another possible explanation for our observations. That is, loci with a relatively high mutation rate could plausibly produce excess rare variants (low TD) and/or excess non-synonymous variation (high *πN/πS*). In this case, we would expect to see the same pattern of category enrichment among genes with extremely high diversity as we do among loci with low TD. We did not, however, observe a consistent enrichment of functional categories among loci with high (or low) values of π or ƟW (S5 Table).

Low values of TD can be observed among loci under selection to remove deleterious mutations. Here, we might expect to observe low levels of non-synonymous variation at the same loci, and perhaps a positive correlation between TD and *πN/πS*. At the between-host scale, there is no overlap between genes in the bottom 5th percentiles of TD and *πN/πS* (Fig. 3). The relationship between TD and *πN/πS* is complex, with a possible switch from a negative to a positive correlation within the distribution. *M.tb* genes with low TD are associated with a range of *πN/πS* values. Ratios of non-synonymous to synonymous variation are an imperfect measure of selection strength for within-population (as opposed to between-species) comparisons [41]: values may be flat over a range of selection coefficients. It is possible that TD and *πN/πS* are responsive to purifying selection in different ways – e.g. over different timescales – and that this is the reason for the non-linear relationship we observed between the two statistics.

**Fig. 3.**
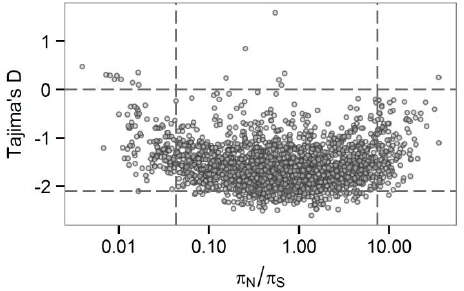
Gene-wise estimates of Tajima’s D and *πN/πS* at the between-host scale. Each circle represents a gene in the H37Rv genome. *π_N_/π_S_* values are plotted on a logarithmic (base 2) scale.

LIP and COG:Q categories contain several large genes, so we wondered about an effect of gene size on TD. LIP and COG:Q categories were enriched for genes in the 95^th^ percentile for size, as were seven other categories (S7 Figure). We observed a weak correlation between TD and gene size (R squared 0.25). While the small number of extremely large genes (>5kb) all have low TD, genes in the 5^th^ percentile of TD have a range of sizes (S8 Figure). One possible explanation for an association between gene size and TD is an effect of the target size for deleterious mutations, with larger genes providing larger mutational targets. Given the strong linkage of sites in the *M.tb* genome, these effects may be particularly relevant.

Low values of TD can also be associated with selective sweeps, where an advantageous mutation rises quickly in frequency. In recombining organisms, this can create local reductions in TD around the site of selection following the sweep. However, for organisms like *M.tb* in which linkage of sites is thought to be complete (i.e. strictly clonal [42]), all variants, which are linked to the adaptive mutation, should move in parallel with it. Therefore, we would not expect differences among *M.tb* loci in values of TD to be driven by selection for advantageous mutations. There is, however, little theoretical work to guide the interpretation of locus specific variation in TD for clonal organisms. More work is needed to identify conditions under which regional variation in TD may be observed in the setting of complete linkage of sites. Our observations could result from unexplored neutral and/or selective regimes that give a signature of locus specific patterns of TD in a linked genome. It is also possible that linkage of *M.tb* loci is not in fact perfect, and our results reflect unrecognized recombination during TB infection.

### Patterns of variation at drug targets

As expected, alleles at several known targets of anti-TB drugs underwent extreme changes in frequency between serial, within-host samples (FST outliers, Fig. 4 and S9 Table). This is due to the emergence of drug resistance mutations, which increase in frequency in response to selection pressures imposed by drug therapy. Interestingly, several drug targets were in the low tail (≤ 5^th^ percentile) of TD across numerous within-host samples. We defined a cutoff of 5 or more samples as extreme, based on the observation that < 4% of genes met this criterion. Within-host values of TD were in the extreme low tail in 5 or more samples for *gyrA*, *gyrB*, *embB*, and *ethA*.

**Fig. 4.**
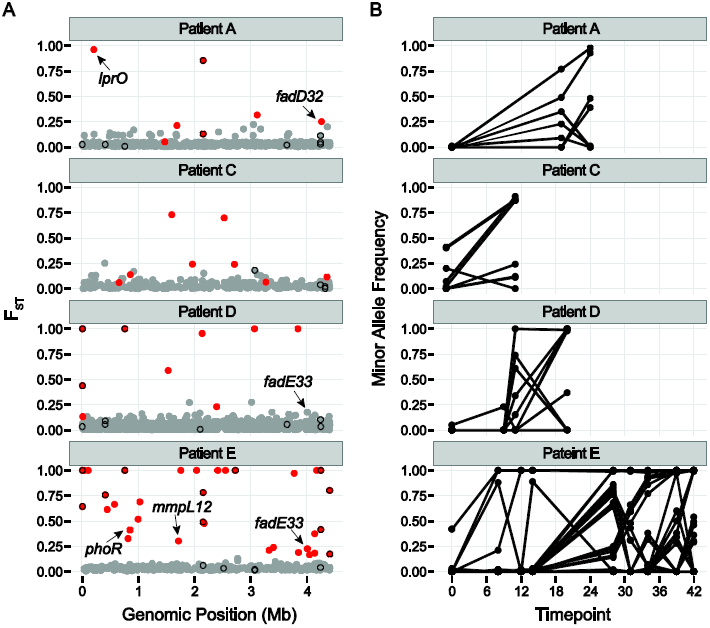
Population differentiation between samples. A.) Pairwise F_ST_ of polymorphic sites. Each patient sample was treated as a population and F_ST_ was calculated individually for each polymorphic site in a population. Calculations were performed on all polymorphic sites covered by at least 10 reads per sample for which the minor allele was supported by a minimum of 6 reads across all samples from the population (*see Methods*). Dots show the highest observed F_ST_ value for each single nucleotide polymorphism (SNP) across the H37Rv genome. Red color coding indicates allele frequencies changed significantly across the sampling interval (based on a Fisher’s exact test; *q*-value < 0.01). Genes implicated in drug resistance are outlined in black. Outliers of the F_ST_ distribution are likely to be under positive selection, or linked to a mutation under positive selection. B.) Allele frequencies over the course of treatment for SNPs with significant changes in allele frequency (red dots in A).

Two recent studies identified *M.tb* genetic variants associated with drug resistance phenotypes using tests for excess polymorphisms and homoplastic polymorphisms, respectively [43,44]. These variants could play a direct role in drug resistance, or they could increase the fitness of bacteria with drug resistance mutations at other loci (compensatory mutations). The ‘targets of independent mutation’ (TIM) category identified by Farhat et al. was enriched among genes with low TD in two of the samples in our study (Fig. 2). Two of the targets identified by Zhang et al, *lprO* and *fadE33*, harbor FST outlier SNPs in our dataset (Fig. 4 and S9 Table). A D->G variant in *lprO* emerged in Patient A’s population and rose to near fixation in parallel with a D->N variant in *katG*. A SNP in *fadE33* emerged in the last sample collected from Patient E. Based on their allele frequencies, it may have been on the same genetic background as a non-synonymous variant in *gid* that is thought to mediate streptomycin resistance. Interestingly, a different SNP in *fadE33* also emerged in Patient D’s *M.tb* population. We were not able to confidently calculate allele frequencies of this SNP across more than one sample, due to low coverage at the site. FadE33’s function is not known; it is essential for *M.tb* growth on cholesterol [45]. Based on these data and the findings of Zhang et al, we hypothesize that LprO and FadE33 play a role in compensation for drug resistance mutations.

### Extreme patterns of variation in mycolic acid synthesis genes

Lipid metabolism genes are found among those with *πN/πS* > 1 at the within-host scale (183/2122 genes with *πN/πS* > 1 are categorized as LIP). In the *M.tb* population of Patient E, a specific sub-group of lipid metabolism genes – those in the mycolate biosynthesis superpathway - are over-represented among genes in the top 5% of values of *πN/πS* at two different time points (E0, *p-*value = 0.026; E12, *p*-value = 0.049), and over-represented among genes in the bottom 5% at one time point (E42, *p*-value 0.031).

Many individual genes in the mycolic acid synthesis pathway exhibited interesting patterns of variation across evolutionary scales. For example, fatty acid synthase (*fas*, *Rv2524c*) is in the 1^st^ percentile of gene-wise TD values in the between-host dataset (BHTD), it was in the low tail of gene-wise TD values for 14 within-host samples (WHTD), and it was one of nine genes with *πN/πS* > 1 across three patients’ *M.tb* populations (Table 2). This gene was also identified as a possible target of positive selection in a recent study of genetic variation among *M.tb* strains in the Beijing lineage [46]. *Pks5*, which is in the mycolate biosynthesis superpathway, was also singled out as a target of selection in Merker et al. *Pks5* was in the 1^st^ percentile of BHTD, it was in the low tail of WHTD in 11 samples, and had *πN/πS* > 1 across three patients’ *M.tb* populations (Table 2).

**Table 2.** Genes with *πNπS* > 1 across 3 within-host *Mycobacterium tuberculosis* populations. Genes identified in at least one sample from 3 independent patients’ *M.tb* population with a *πNπS* > 1 are displayed. Two genes meeting this criteria were excluded due to potential copy number variants identified by manual inspection of the alignments. *Indicates the gene is in the superpathway of mycolate biosynthesis.

| Gene | Common Name | Gene Description |
| --- | --- | --- |
| *Rv0101* | *nrp* | peptide synthetase |
| *Rv0931c* | *pknD* | transmembrane serine/threonine-protein kinase D |
| *Rv1527c* | *pks5* * | polyketide synthase |
| *Rv2318* | *uspC* | periplasmic sugar-binding lipoprotein |
| *Rv2476c* | *gdh* | NAD-dependent glutamate dehydrogenase |
| *Rv2524c* | *fas* * | fatty-acid synthase |
| *Rv2566* |  | conserved hypothetical protein |
| *Rv2946c* | *pks1* * | polyketide synthase |
| *Rv3157* | *nuoM* | NADH dehydrogenase I chain M |

FadD32 is essential for mycolic acid synthesis [47] and is currently being investigated as a target for new TB treatments [48,49]. Patient A’s *M.tb* population harbored an FST outlier SNP in *fadD32* (Fig. 4 and S9 Table). Expression of *fadD32* was recently found to decrease over the course of infection in a patient with extensively drug-resistant TB [25]. Eldolm et al speculated that down-regulation of *fadD32* could help to compensate for drug resistance mutations. In addition to identifying this gene in our outlier FST analysis, we found that it had one of the highest gene-wise values of *πN/πS* in the between-host sample (*πN/πS* = 32), as well as being in the 93^rd^ percentile of BHTD. These results suggest that FadD32 is a target of selection during infection and possibly transmission, and that this leaves a signature of diversifying selection at the between-host scale. Although regulatory and/or sequence variation at this locus could play a role in compensation for drug resistance mutations, we think it is unlikely that this alone would produce the extraordinary levels of diversity observed in our study. Despite recent pre-clinical promise of FadD32 as a target of coumarin compounds [48,49], our finding of high diversity in this gene suggests that the genetic barrier to acquisition of resistance at this locus is likely to be low.

The gene encoding phosphopantetheinyl transferase (*pptT*) harbored a non-synonymous SNP at a frequency of 41% in the *M.tb* population of patient B, which was undetected in the prior sputum sample (data from this sample did not meet inclusion criteria). *PptT* belongs to the functional categories LIP and COG:Q. PptT activates Pks13, a type-I polyketide synthase involved in the final step of mycolic acid biosynthesis, as well as various type-I polyketide synthases required for the synthesis of lipid virulence factors [50,51]. *Pks13* was found to be down-regulated during infection in the study of Eldholm et al. Active FadD32 and Pks13 are involved in the final steps of mycolic acid condensation *in vitro* [47]. The identification of an F_ST_ outlier in *fadD32* (patient A) and a newly emergent SNP in *pptT* (patient B), the association of mycolic acid synthesis genes with extreme values of TD and *πN/πS*, and the identification of these genes as targets of selection/regulatory variation across independent studies suggests that *M.tb* mycolic acids may be modulated over the course of individual TB infections. This individual-to-individual variation could explain the high diversity observed at the between-host scale in some of these genes. Caution is warranted in drug development aimed at this pathway, as loci with high natural diversity may not be optimal drug targets.

### Variation in polyketide synthases and related genes

*M.tb* polyketide synthases (PKS) play essential roles in the biosynthesis of lipids and glycolipids of the cell envelope [52]. These lipids and glycolipids are positioned at the outer edge of the envelope, at the host-pathogen interface. As might be predicted based on their location, they play important roles in the pathogenesis of TB (reviewed in [52–54]). In addition to mycolic acids, PKS-synthesized lipids include phthiocerol dimycocerosates (PDIM), sulfolipids (SL), polyacyl trehalose (PAT), diacyl trehalose (DAT), phenolic glycolipid (PGL), novel complex polar lipids synthesized by Pks6 (POL), and mannosyl-β-1-phosphomycoketides (MPM).

Genes encoding polyketide synthases (PKS) are particularly striking for extreme patterns of variation: among 22 *PKS* homologs in the H37Rv genome (excluding the incorrectly annotated *pks16*, and counting *pks15/1* and *pks3/4* as one locus each [52]), we found 18 exhibited extreme patterns of variation (5^th^ or 95^th^ percentile at the between-host scale, and/or in the extreme low tail of WHTD in ≥ 5 samples). The most common pattern found among *PKS* is low gene-wise TD at both within- and between-host scales. Of 15 genes found in the 5^th^ percentile of BHTD and the low tail of WHTD in ≥ 10 samples, 10 are PKS.

Similar to *pks5* (discussed above), *pks1* is also among nine genes with a *πN/πS* > 1 across three patients’ *M.tb* populations(Table 2). Zhang et al [43] found evidence of an association between *pks2*, *pks8* and *pks17* variants and drug resistance. *Pks2* and *pks8* exhibited extreme patterns of variation in our study (1^st^ percentile of BHTD and low tail of WHTD in 15 samples; 1^st^ percentile of BHTD and low tail of WHTD in 11 samples). Farhat et al [44] found evidence of positive selection on *ppsA*, *pks3* and *pks12*, all of which were in the extremes of BHTD and/or 5 or more samples’ WHTD in our data.

PKS-synthesized lipids are transported to the cell surface by the Mycobacterial membrane protein Large (MmpL) family of membrane proteins. H37Rv includes genes encoding 13 MmpL transporters [55]; substrates have not been identified for all them. MmpL have been shown to play important roles in TB pathogenesis [55,56]. *MmpL* genes were also notable for extreme patterns of variation: 10/13 of these loci were in the extreme tail of *πN/πS* and/or TD at the between-host scale and/or the low tail of TD in 5 or more within-host samples. Similar to *PKS* genes, the combination of low values of gene-wise TD at the within- and between-host scales was most common. In addition, we identified an FST outlier SNP in *mmpL12* (Fig. 4 and S9 Table); *mmpL11* has been identified as a target of positive selection in a previous study [46] and *mmpL1* variants were associated with drug resistance in a separate study [43]. Genes encoding two non-MmpL transporters of PDIM, *drrA* and *drrC*, also exhibited extreme patterns of variation. *DrrA* is in the 99^th^ percentile of BHTD, whereas *drrC* is in the 1^st^ percentile of between-host *πN/πS.* These observations suggest that PKS and associated transporter genes are involved in adaptation of *M.tb* during infection. For some of these loci, there is evidence suggesting a role in drug resistance, augmenting the phenotype and/or increasing the fitness of drug resistant mutants.

### Patterns of variation among regulators of PKS lipids

We observed extreme patterns of variation in genes involved in regulation of immunomodulatory lipids, in addition to those involved in their biosynthesis and transport (Fig. 5). For example, the gene encoding a regulator of PDIM synthesis, *pknD*, was in in the low tail of BHTD and six samples’ WHTD. It was also in the group of nine genes with *πN/πS* > 1 across three patients’ *M.tb* populations (Table 2). PknD is thought to regulate deposition of PDIM on the cell wall via its effects on MmpL7, which transports PDIM [57]. Another example is the GTP pyrophosphokinase *relA* (*Rv2583c*), which regulates *M.tb* gene expression during the chronic phase of murine infection and plays a central role in the response to hypoxia and starvation [58,59]. *RelA* was in the low tail of seven samples’ WHTD. RelA modulates expression of *pks3/4*, *pks5* and *fas*, all of which exhibited extreme patterns of variation (discussed above).

**Fig. 5.**
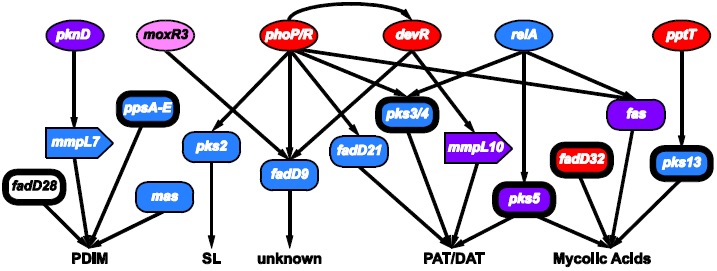
Connectivity of regulators and targets involved in the synthesis of *Mycobacterium tuberculosis* immunomodulatory lipids. Interactions (shown as arrows) among select genes involved in the regulation (circles), synthesis (rectangles) and transport (block arrows) of phthiocerol dimycocerosates (PDIM), sulfolipids (SL), polyacyl trehalose (PAT), diacyl trehalose (DAT), and mycolic acids. The coloring scheme reflects patterns of variation observed at these loci: genes colored red had signatures of positive selection (gene contained an F_ST_ outlier and/or was in the 95^th^ percentile of *πN/πS* at the between-host scale), blue indicates the gene had extreme Tajima’s D (TD) (5^th^ percentile of between-host TD and/or 5^th^ percentile of within-host TD for ≥ 5 within-host samples), purple genes contained both low TD & signatures of positive selection, while pink signifies the gene had evidence of selection against non-synonymous diversity (5^th^ percentile between-host *πN/πS*). Genes identified as targets of selection in other studies are indicated with bolded outlines. See discussion in text for more details.

Patterns of variation in *fadD9*, which is of unknown function, are similar to many genes affecting immunomodulatory lipids (S10 Table): *fadD9* was in the 2^nd^ percentile of BHTD and the low tail of 10 samples’ WHTD. Expression of *fadD9* is under the control of two regulators that are central to *M.tb*’s adaptation to the within-host environment: PhoP [60] and DevR [61] (Fig. 5). PhoP controls the production of PAT/DAT and SL [62]. *DevR* exhibited extreme patterns of variation, with BHTD in the 99^th^ percentile. Another regulator of *fadD9*, *moxR3*, exhibited extreme patterns of variation: this gene was in the 5^th^ percentile of between-host πN/πS (Fig. 5 and S10 Table). *MoxR3* is upregulated during re-aeration of hypoxic *M.tb* cultures [63]. As noted above, the function of FadD9 has not been identified; patterns of variation in this gene and its regulators, as well as its co-regulation with virulence associated lipids, suggest it may play a role in host-pathogen interactions, and that it is a target of selection within hosts.

### PKS lipids as targets of selection within hosts

We have shown that numerous genes affecting PKS associated lipids (mycolic acids, PDIM, PAT, DAT, PGL, POL, and MPM) exhibit extreme patterns of variation across a range of statistical measures and independent data sets. We hypothesize that these patterns of variation are due to adaptation within *M.tb*’s natural human host. Several characteristics of PKS-associated lipids make them likely targets of selection during infection.

PKS associated lipids are important mediators of host-pathogen interactions. For example, mycolic acids are known to modify host cell phenotype [64] and to affect virulence in animal models of TB [65]. PDIM has major impacts on *M.tb*’s virulence in animal models of TB [66,67] via numerous mechanisms (reviewed in [68]) including protection from innate immune responses [69]. Like PDIM, the PKS-synthesized lipids PAT/DAT, SL, MPM, POL and PGL also appear to mediate interactions between *M.tb* and the immune system [66, 70–86].

The capacity for adaptation is predicated on phenotypic variation, which has been demonstrated for PKS associated lipids. Production of PAT/DAT has been shown to vary among clinical isolates; some isolates do not make them at all [87]. Production of PGL also varies among clinical isolates, and variant forms of PDIM have been identified in clinical isolates [54]. Based on their roles in pathogenesis and their variability, it has been proposed previously that modifications of these lipids could arise during natural infection in order to optimize host cell manipulation [54]. The observation of unusual patterns of variation at genes involved in the synthesis, transportation and regulation of these lipids provides evidence in support of this hypothesis.

Several features are conducive to efficient selection on genes controlling the synthesis, regulation, and transportation of PKS associated lipids. There are connections between biosynthetic pathways controlling production of distinct PKS synthesized lipids (e.g. PDIM, SL, and mycolic acids) such that metabolites can be shuttled between them [88,89]. This flexibility allows *M.tb* to respond rapidly to environmental fluctuations. It may also allow more efficient selection for adaptive mutations, since single mutations can potentially affect multiple lipid products. In addition, since intermediate metabolites could potentially be shuttled down multiple pathways, they need not accumulate with potentially toxic effects if one pathway is affected by a harmful mutation.

PDIM, PAT/DAT and SL are not essential for growth of *M.tb* in artificial media, and PDIM is in fact frequently lost during passage of *M.tb* in the laboratory [87,90]. This suggests that functions of these lipids are specific to natural, within-host environments. The expression of *M.tb* immunomodulatory lipids is responsive to physiological conditions (hypoxia, starvation) encountered during natural infection [91,92]. In tissue culture and animal models of TB, the shift from axenic growth to infection is accompanied by alterations in expression of genes involved in lipid metabolism with consequent changes to the bacterial cell envelope [88,93–95]. Further supporting the idea that these lipids are important for adaptation to the pathogenic niche, MPM, PGL, and PDIM are only found among pathogenic mycobacteria, and DAT/PAT and SL are specific to the *Mycobacterium tuberculosis* complex (MTBC) (Fig. 6) [54,62,79]. It was shown recently that even among members of the MTBC, production of DAT/PAT and SL is specific to human-pathogenic mycobacteria [62].

**Fig. 6.**
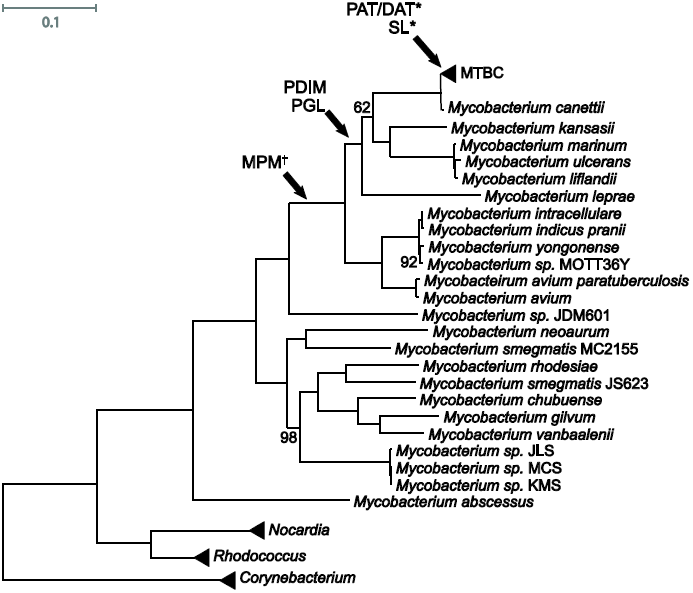
Mycobacteria maximum likelihood tree. Phylogenetic analysis is based on a core genome alignment of 57 strains (S9 Table). Arrows indicate the most probable emergence of specific lipids in the phylogenetic history of Mycobacteria based on previous studies [54,62,79]. *Indicates lipids are only found in *M. tuberculosis sensu stricto*, not other members of *Mycobacterium tuberculosis* complex (MTBC). †Indicates uncertainty in the placement because lipids of *M. sp.* JDM501 have not been characterized. Bootstrap values are 100 unless otherwise labeled. Scale bar indicates the mean number of substitutions per site. ABREVIATIONS: PDIM – phthiocerol dimycocerosates; PGL – phenolic glycolipids; SL – sulfolipids; MPM – mannosyl-β-1-phosphomycoketides; PAT – polyacyl-trehalose; DAT – diacyl-trehalose

Recent work has shown that PAT/DAT, PDIM, and SL all impair phagosomal acidification and thereby improve *M.tb* survival within macrophages; there appears to be significant flexibility in how these functions are performed, and distinct lipid moieties may compensate for each other [87]. By providing cover during exploration of the fitness landscape, both functional redundancy and metabolic flexibility may increase the potential for rapid adaptation. For example, they would enable progress toward fitness peaks that are otherwise unreachable as a result of a preceding ‘valley’ created by mutations that are deleterious except in certain combinations. This may be particularly important for a clonally reproducing organism such as *M.tb*.

## Conclusions

We have shown that genetic diversity of *M.tb* populations can vary dramatically between individuals with TB. Variation correlated with clinical severity, rather than treatment status, drug resistance phenotype or *M.tb* lineage. Further studies are needed to explore the replicability of patterns observed here across different types of patients with TB and treatment settings.

We have also observed distinct patterns of variation among genes associated with *M.tb* virulence lipids: these patterns carry through functional links among genes, and across different statistical measures and datasets in our own and previously published studies. These lipids are good candidates for adaptation during infection, based on the phenotypic variation observed among clinical isolates, their roles mediating host interactions, their specific association with the pathogenic niche, and their flexible functional architecture However, it is important to note uncertainties in the data and its interpretation. We cannot, for example, assert that all genetic variants described here were generated within hosts. In all the studies whose data we analyzed, clinical specimens were manipulated *in vitro* prior to *M.tb* genome sequencing. Some mutations may have occurred during culture.

Genetic variation in *M.tb* is skewed to rare allele frequencies, at both the within- and between-host scale. While we have endeavored to ameliorate the impacts of spurious variants introduced by sequencing errors and errors introduced during processing of sequencing read data, rare variants are difficult to disentangle from sequencing errors, particularly in pooled sequencing data. As is standard in studies of *M.tb* genomics, our analyses excluded genomic segments (totaling ~11%, see *Methods*) that are poorly resolved with short read sequencing technologies. Additional measures to reduce the effects of sequencing errors included trimming low quality bases from sequencing reads, implementing mapping and base quality thresholds, and excluding indels and SNPs demonstrating strand-bias and/or tail-distance-bias. We used PoPoolation, which explicitly accounts for the effects of sequencing error, to estimate population genetic parameters in our study. Validation studies of PoPoolation have shown that implementation of PoPoolation’s recommended minor allele count and quality thresholds decreases the error rate from 1% to ~ 0.01% post processing [27]. These studies also demonstrated that PoPoolation parameter estimates from pooled sequencing data are highly correlated with estimates from Sanger sequencing data.

However, there are differences between PoPoolation validation datasets and those analyzed in our study: PoPoolation was validated for sequencing reads generated on the Illumina GAIIx from a *Drosophila melanogaster* population (and data simulated to mimic these reads), whereas we have analyzed sequencing reads generated on the Illumina HiSeq from *M.tb* populations. GAIIx and HiSeq differ in their biases [96], and GC content differs significantly between *M.tb* and *D. melanogaster*. We note that despite these potential limitations of PoPoolation in correcting errors in our pooled data, we observed similar patterns of genetic variation in the between-host dataset, which is based on single-colony isolates and a different variant-discovery pipeline.

Interpretation of genetic data across evolutionary scales is also not straightforward. The consistent identification of specific functional categories in extremes of various statistics suggests that they are due to natural selection, but the type of selection and where it operates is not clear. The patterns we observed may be due to selection imposed by drug therapy, the immune response and/or transmission, with further modification as a result of complex neutral influences on variation. In addition, our hypotheses need to be investigated at the level of phenotypes. Based on the analyses presented here and their contextualization with published data, we propose that further characterization of individual-level, phenotypic variation in *M.tb* PKS associated lipids is likely to be fruitful. In addition, several of the loci described here are worthy of investigation for a potential role mediating or compensating for drug resistance.

## Methods

### Data Collection for Within- and Between-Host Datasets

**Within-host *M.tb* data set (*n* = 19)**. We used carefully chosen WGS data from three previously published studies [23,25,26] to characterize within-host populations of *M.tb*. In each of these studies, primary specimen from sputum samples of patients being treated for TB were subcultured on Lowenstein-Jensen (LJ) slants without single colony passage; genomic DNA was extracted from each LJ slant and sequenced on an Illumina platform to capture the *M. tb* population present in the sputum sample (pool-seq). Since sequencing error biases are known to vary across platforms [96], we only used WGS data generated on the Illumina HiSeq (data from a variety of platforms were analyzed in the original Sun et al and Eldholm et al studies). As low frequency variants are important in the analysis of population genetic parameters, we only used samples for which the mean coverage across the genome was ≥ 50X in the present study. Accession numbers and publicly available meta information for the 19 within-host samples (from 5 different patients’ *M.tb* populations) passing these criteria are shown in Table 1. In each of the primary studies, standard genotyping methods (restriction fragment length polymorphism analysis and/or variable number tandem repeat analysis) indicated that all serial isolates had identical profiles. Accession numbers, mapping statistics, and exclusion basis (when applicable) for all sequencing runs considered are in Table S1.

**Between-host *M.tb* data set (*n* = 201).** We used previously published WGS data from 201 diverse, globally extant strains of *M.tb* [29] to characterize between-host populations of *M.tb*. The data set includes isolates from all seven major lineages of *M.tb* [97,98]. Accession numbers and more detailed information about the strains are in S2 Table.

### Processing of raw sequencing reads

For the within-host *M.tb* data set, we trimmed low-quality bases from the FASTQ data using a threshold quality score of 20, and reads of length less than 35bp were discarded using Trim Galore! (http://www.bioinformatics.babraham.ac.uk/projects/trim_galore/) - a wrapper tool around Cutadapt [99] and FastQC (http://www.bioinformatics.babraham.ac.uk/projects/fastqc/). We mapped reads to H37Rv (NC_000962.3) [100] using the default parameters of the BWA MEM algorithm with the –M flag enabled for downstream compatibility [101], and we removed duplicates using Picard Tools (http://picard.sourceforge.net). Local realignment was performed with the Genome Analysis ToolKit (GATK) [102] in a sample aware manner, and aligned reads with mapping quality > 20 were converted to mpileup format using Samtools software [103].

The resulting reference-guided assembly of each sample spanned over 98.9% of the H37Rv genome, with a mean depth of coverage per site ranging from 56X to 192X (S1 Table). Loci at which indels were present in a sample were removed along with 5bp of flanking sequence using the PoPoolation package [27].

For the between-host *M.tb* data set, we trimmed low-quality bases from FASTQ data using a threshold quality of 15, and reads resulting in less than 20bp length were discarded using Trim Galore! (http://www.bioinformatics.babraham.ac.uk/projects/trim_galore/) - a wrapper tool around Cutadapt [99] and FastQC (http://www.bioinformatics.babraham.ac.uk/projects/fastqc/). Reads were mapped to H37Rv (NC_000962.2) [100] using the suggested algorithm of BWA for the given sequencing strategy (e.g. paired-end/single-end, read length) [101,104], and duplicates were removed using Picard Tools (http://picard.sourceforge.net). We used GATK to perform local realignment and variant calls using a minimum phred-scaled confidence threshold of 20 [102]. Variants were filtered with the following expression “QD < 2.0 || FS > 60.0 || MQ < 40.0 || MQRankSum < −12.5 || ReadPosRankSum < −8.0” as described in the GATK best practices. Genome alignments were generated with scripts that can be found at https://github.com/tracysmith/RGAPepPipe.

Transposable elements, phage elements, and repetitive families of genes (PE, PPE, and PE-PGRS gene families) that are poorly resolved with short read sequencing were removed from the mpileup files and alignment prior to subsequent analyses plus 5bp up- and down-stream of the genes. We additionally removed regions that were found to have poor mapping quality using the CallableLoci tool of the GATK: for each within-host sample, any region reported as poorly mapped using the following flags was removed from all datasets (including between-host) plus 5bp up- and down-stream: -frlmq 0.04 –mmq 20 -mlmq 19. As an additional measure, we also removed sites with the lowest 2% average mapping quality in the global alignment from all datasets prior to subsequent analysis. Regions removed from all datasets based on these criteria can be found in S12 Table. Polymorphisms in the within-host dataset demonstrating strand-bias or tail-distance-bias identified by a previously described method [22] (https://github.com/tamilieberman/IntrasamplePolymorphismCaller) were removed along with 5bp up- and down-stream from all within-host samples (S12 Table).

### Population Genetic Estimates of Within-host Populations

We used the PoPoolation package [27] to estimate nucleotide diversity (π), Watterson’s theta (ƟW), and Tajima’s D (TD) in sliding windows across the *M.tb* genome. Sensitivity analyses of these statistics to input parameters are described in the S3 File. Parameter estimates were strongly influenced by sequencing coverage (S3 File). In order to alleviate biases in parameter estimates caused by variable coverage among samples, we randomly sub-sampled read data without replacement to a uniform coverage of 50X; this process was performed 9 times for each sample to reduce potential biases introduced by sampling of rare alleles. We used default equations in the PoPoolation package to estimate π, ƟW, and TD [27,28]. In order to reduce the effects of sequencing errors on parameter estimates, PoPoolation implements modified calculations of the classical π, ƟW, and TD that are only evaluated on SNPs above a designated minor allele count. PoPoolation estimators account for the truncated allele frequency spectrum and re-sequencing of the same chromosomes that occurs during pool-seq. PoPoolation’s recommended minor allele count and quality thresholds are based on simulation studies indicating that their implementation decreases the error rate of 1% in the raw data to ~0.01% in the processed data. PoPoolation parameter estimates from pooled data have also been found highly concordant with estimates from Sanger sequencing data.

Unless otherwise noted, all calculations performed with the PoPoolation package were implemented using the recommended minimum minor allele count of 2 for 50X coverage, and a pool-size of 10,000 (*see S3 File for justification*). Calculations performed under differing parameters can be found in the Supplemental Data. Sliding-window analyses were performed using a window-size of 100K and a step-size of 10K; data presented are the mean of each window across 9 replicate sub-sampled mpileups (Fig. 1). Genome-wide averages are expressed as the mean across all windows sufficiently covered (≥ 60% coverage of 50X) (S4 Table).

To calculate gene-by-gene estimates of π, ϴW, and TD from the 50X sub-sampled mpileups, we again used the PoPoolation package [27]. Gene coordinates were obtained from the standardized Gene Transfer Format (.gtf) for the H37Rv annotation on the TB database (tbdb.org) (S13 file) [105]. Excluding genes with inadequate coverage (<50% of the gene), we calculated the mean value of each statistic across the 9 replicate sub-samplings for each gene of each sample, and compared it to all other genes within the sample.

Using the same .gtf file and the PoPoolation package, we also calculated the average number of nonsynonymous differences per nonsynonymous site (*πN*) and the average number of synonymous differences per synonymous site (*πS*) for each gene in the 50X sub-sampled mpileups. Recognizing that chance samplings of very rare mutations in one replicate sub-sampled mpileup would lead to skewed distributions, we took the median values of *πN* and *πS* across 9 replicate sub-samplings for each gene of each sample and calculated *πN/πS*. Excluding genes with inadequate coverage (<50% of the gene) and genes with *πN* and/or *πS* equal to zero, *πN/πS* values were compared relative to all genes passing these criteria within the sample.

To identify candidate SNPs under selection, we treated each temporal sample of a patient as a population, and estimated pairwise FST for each variable site in the genome with PoPoolation2 [37].. Unlike the sliding-window and gene-wise estimates, all sequencing data for which the minimum base quality was ≥ 20 was considered (i.e. no sub-sampling). We excluded all sites with a coverage < 10, and only considered those sites with a minimum allele count ≥ 6 (pooled across all samples for a patient). F_ST_ estimates were subjected to an empirical outlier analysis. To reduce biases resulting from variable coverage, we conditioned our analysis on the ability to detect a significant change in allele frequency between samplings using a two-sided Fisher’s exact test as previously proposed [38]. To account for the large number of tests performed we used a false discovery rate (FDR) of 5% and calculated adjusted *p*-values (*q*-values) (R stats Package [106]). SNPs with extreme FST values (> 0.1) and a *q*-value < 0.01 were deemed outliers and are listed in S9 Table.

S14 Table shows SNPs passing all filtering criteria that were used in the above described population genetic estimates for the within-host populations.

### Population Genetics of Globally Extant Strains (Between-Host)

We used PoPoolation [27] with the “disable-corrections” flag enabled (calculations are performed using the classical equations) to generate sliding-window estimates of π and ƟW from whole genome alignments of all 201 globally extant *M.tb* strains, *M.tb* isolates from lineage 2 (East Asian lineage, *n* = 37), and *M.tb* isolates from lineage 4 (Euro-American lineage, *n* = 53) (S2 Table) [29]. We required a minimum of 75% of the strains have non-missing data for a site to be included in the analysis (“minimum coverage”), set the “minimum minor allele count” to one, and the “pool-size” to the number of strains being analyzed.

We calculated gene-by-gene estimates of π, ϴ, TD, πN, and πS for the global dataset in the same way as was done for within-host samples, save for a few exceptions: rather than use the default PoPoolation estimators that apply corrections for sequencing errors [27,28], we employed the “disable-corrections” flag (calculations are performed using the classical equations), set the “minimum minor allele count” to one, and the “pool-size” was set to 201 to reflect the number of strains in the dataset; the “minimum coverage” was set to 151 to exclude any genes that were not represented by at least 75% of strains in the dataset. Finally, no averaging was performed. For *πN/πS* calculations, we again excluded genes with *πN* and/or *πS* equal to zero, and compared values relative to all genes passing these criteria within the sample.

### Functional & Pathway Enrichment Analyses

For each within-host sample, between 58.7–90.1% of annotated genes in the H37Rv genome had sufficient coverage to calculate TD; for the between-host sample, TD was calculated for 90.2% of annotated genes in the H37Rv genome, as this was the fraction covered by at least 75% of the strains. Genes with TD values in the top and bottom 5% of the distribution for a given sample were deemed candidate genes of selection. The significance of enrichment for functional categories in candidate genes of selection was assessed with a two-sided Fisher’s exact test. To account for multiple hypothesis testing, we used a false discovery rate of 5% and calculated *q*-values (R stats Package, [106]). We used the following annotation categories to classify *M.tb* genes: computationally predicted Clusters of Orthologous Groups (COG) (*n* = 21 categories) [40]; essential and nonessential genes for growth *in vitro* as determined by transposon site hybridization (TraSH) mutagenesis [107]; genes essential for growth in a murine model of TB [108]; "targets of independent mutation" associated with drug resistance [44]; and the *M.tb*-specific, manually curated functional annotation lists from TubercuList (*n* = 7) [39]. COG annotations for the H37Rv genome were obtained from the TB database (tbdb.org) [105], as were the TraSH “*in vitro* essential”, “*in vivo* essential”, and “nonessential” gene annotations. TubercuList functional annotations were obtained from tuberculist.epfl.ch [39] and reflect the most up-to-date annotations when the database was accessed (12/01/2013). We did not include the TubercuList categories “PE/PPE”, “stable RNA”, and “unknown” in our analyses. S15 File contains the genes included in each category. Functional enrichment of genes in the top 5% of π and ϴW for each sample were performed in the same manner.

Variable proportions of annotated genes in the H37Rv genome had sufficient coverage and contained both nonsynonymous and synonymous variation in a given within-host sample. While a large percentage of genes were also excluded from the analysis because they lacked synonymous and/or non-synonymous variation, the inference to be drawn from an elevated *πN/πS* value still holds. To look for commonalities among patients, we examined all genes found to have a positive *πN/πS* value in at least one sample from multiple patients; genes found to have a *πN/πS* > 1 in three patient *M.tb* populations are shown in Table 2. Upon noticing a preponderance of genes in the superpathway of mycolate biosynthesis on the cellular overview tool of the TB database (tbdb.org) [105] we formally tested for an enrichment of genes in this category using a two-sided Fisher’s exact test.

### Methods for variant quality & mapping quality

We used the Python package pysamstats (https://github.com/alimanfoo/pysamstats) to calculate the root-mean-square (RMS) value of base qualities for variant alleles and the RMS mapping quality for reads aligned at such polymorphic sites in the reference-guided-assemblies of within-host samples. Variants occurring in genes categorized as CHP, LIP, or COG:Q were subject to a one-sided Student’s T-test to determine whether the mean RMS base qualities of the category was significantly lower than that occurring in any gene.

### Mycobacteria Core Genome Alignment and Phylogenetic Tree

We downloaded finished genomes of mycobacterial species from NCBI (S11 Table). Reference guided assemblies for species where only sequencing reads were available were performed as described above for the between-host dataset. We used Prokka v 1.7 [109] for genome annotation. Protein sequences output by Prokka were clustered into orthologous groups using OrthoMCL [110]. The core proteins (those found only once in every genome) were aligned using MAFFT [111], trimmed with trimAl [112], and concatenated. Scripts used for core genome analysis can be found at https://github.com/tatumdmortimer/core-genome-alignment. We used RAxML v 8 [113] for maximum likelihood phylogenetic analysis of the core alignment. We used Dendroscope [114] for tree viewing and editing.

#### Acknowledgements

We thank Jennifer Bratburd for her contribution to the generation of the Mycobacteria species tree, Tracy M. Smith for her input on the manuscript, and Karl Broman for useful input on the statistical analyses and data visualization.

## Supplementary Files

**S1 Table. Summary statistics and meta information for within-host samples considered for this study.** We used carefully chosen whole genome sequence data from three previously published studies [23,25,26] to characterize within-host populations of *M.tb*. In each of these studies, primary specimen from decontaminated sputum samples of patients being treated for TB were sub-cultured on Lowenstein-Jensen slants without single colony passage; genomic DNA was extracted from each slant and sequenced on an Illumina platform to capture the *M. tuberculosis* population present in each sample (pool-seq). Inclusion criteria for this study was threefold: 1.) We only used WGS data generated on the Illumina Hi-Seq platform. 2.) We only used samples for which the mean depth of coverage was > 50X. 3.) Only one sequencing run per sample was used to avoid biases introduced by combining data across multiple runs. Tab A contains information pertaining to all strains considered including sample names, accession numbers, and exclusion criteria where applicable. Tab B contains additional base quality and mapping statistics for samples selected for use in the current study.

**S2 Table. *Mycobacterial tuberculosis* strains used for global and regional datasets in this study.** Strains used in this study are a subset of those used in a previous study [29]. Information pertaining to the place of birth of the patient, the place of isolation of the strain, and the phylogeographic area are taken from Comas et al. - Supplementary Table 1 and reported here for the ease of the reader. We performed phylogenetic analysis on the selected strains and confirmed the lineages reported by Comas et al. Accession numbers are listed for each strain.

**S3 File. Sensitivity analysis and parameter choice justification for PoPoolation software.** Supplementary information on our sensitivity analysis of the PoPoolation Software [27]. This document includes justification of our parameter choices, as well as four figures with legends in the document.

**S4 Table. Genome-wide estimates of nucleotide diversity (π) and Watterson’s theta (ϴW) under varying parameters.** Reference-guided assemblies for within-host samples were subsampled without replacement to a uniform coverage of 50X. Sliding-window analyses of π and ϴW were performed with default PoPoolation equations that account for sequencing error in pooled-data [27]. A window-size of 100Kb, a step-size of 10Kb, and a “pool-size” of 10,000 were used (*see S3 File for justification*). Estimates for within-host samples were generated under three parameter sets: mbq20 – a “minimum base quality score” of 20 with a “minimum minor allele count” of 2; mbq30 – a “minimum base quality score” of 30 with a “minimum minor allele count” of 2; mc3 – a “minimum base quality score” of 20 with a “minimum minor allele count” of 3. Genome-wide estimates are expressed as the mean across all windows covered by at least 60% under the given parameters. Genome-wide estimates for the global and lineage-specific datasets were performed with classical equations in PoPoolation (“disable corrections” flag enabled). Only sites covered by >75% of strains were included in the analyses. A window-size of 100Kb and a step-size of 10Kb were used, and genome-wide estimates are expressed as the mean across all windows passing criteria. Other parameters were not applicable to between-host datasets. 32

**S5 Table. Functional enrichment analysis of genes with extreme values of Tajima’s D (TD), nucleotide diversity (π), and Watterson’s theta (ϴW).** Genes with TD, π, and ϴW values in the top and bottom 5% of the distribution of each sample were tested for enrichment of functional categories (described in *methods*) using a two-sided Fisher’s exact test. To account for multiple hypotheses testing, a false discovery rate of 5% was used and the resulting *q*-values are reported. Red font and cell highlighting indicates significance at the 0.05 level. Note that the results presented in the tab “TD-bot5%” are visualized in Fig. 2.

**S6 Figure. Distributions of base and mapping quality scores of polymorphic sites in notable categories.** Box-and-whisker plots of (A) the root-mean-square (RMS) value of base qualities for variant alleles and (B) the RMS mapping quality for reads aligned at polymorphic sites in the reference-guided-assemblies (pooled across all within-host samples). RMS values were calculated with the python package pysamstats https://github.com/alimanfoo/pysamstats. Distributions are shown for polymorphisms occurring in any gene (black), TubercuList “conserved hypotheticals” (blue), TubercuList “lipid metabolism” (red), and COG:Q “secondary metabolites biosynthesis, transport, and catabolism” (purple). Upper and lower whiskers delineate highest values within 1.5 times the distance between the first and third quartiles. Outliers and plotted as points.

**S7 Figure. Enrichment of functional annotation categories among genes in the 95^th^ or greater percentile for gene length.** The total gene length covered by at least 75% of between-host strains were compared, and the top 5% of genes in the distribution were tested for overrepresentation of functional categories using a two-sided Fisher’s exact test. Genes which were not covered by at least 75% of strains for more than half of the total gene length were excluded.

**S8 Figure. Between-host Tajima’s D (TD) versus gene-length.** Each point corresponds to a gene in the H37Rv genome. Between-host, gene-wise values of TD are plotted against the length of the gene that was at sufficient coverage for the calculation. Genes for which less than half of the gene was covered by 75% of the strains have been excluded. The black dotted line marks the 5^th^ percentile of TD.

**S9 Table. SNPs with extreme FST values in serial samples of within-host *Mycobacterium tuberculosis* populations.** SNPs found to have an FST > 10 and a *q*-value ≤ 0.01 (*see Methods*) are annotated for each Patient. Allele frequency change between longitudinal samples, nucleotide change, and amino acid change are with respect to the minor allele of the first sample time point of each patient (not the H37Rv reference). FST values reported for Patients A, D, and E are the maximum observed value among all possible pairwise comparisons.

**S10 Table. Gene-by-gene estimates of population genetic parameters from within- and between-host samples.** For each gene in the H37Rv genome, population genetic parameters estimated for global (g) and within-host (accession no.) samples are displayed; column headers correspond to the sample followed by one of the following abbreviations: cov – fraction of gene resolved at sufficient coverage; theta – Watterson’s theta (ϴW); pi – nucleotide diversity (π), TD – Tajima’s D; piN – the number of nonsynonymous mutations per nonsynonymous site (*πN*); piS – the number of synonymous mutations per synonymous site (*πS*); piNpiS - (*πN/πS*). Genes had to 33 be resolved at ≥ 50% (and in at least 75% of strains for the between-host dataset) in order for statistics to be calculated. See *methods* for details of how each parameter was calculated. Gene symbols, common names, and descriptions are from S12 File obtained from tbdb.org. Calculations for within-host samples were calculated with PoPoolation [27] under three different parameter sets and are presented under different tabs: mbq20 – a “minimum base quality score” of 20 with a “minimum minor allele count” of 2; mbq30 – a “minimum base quality score” of 30 with a “minimum minor allele count” of 2; mbq20_mc3 – a “minimum base quality score” of 20 with a “minimum minor allele count” of 3.

**S11 Table. Strain names and accession numbers for genomes used in Figure 7.** Strain names and accession numbers are listed for the genomes used to generate the core genome alignment and maximum likelihood tree.

**S12 File. Regions removed from reference-guided assemblies.** Stringent quality filters were imposed in our data processing pipeline. A) Non-overlapping regions removed from all datasets (within-host and between-host): transposable elements, phage elements, and repetitive families of genes (PE, PPE, and PE-PGRS gene families) that are poorly resolved with short read sequencing, regions found to have poor mapping quality using the CallableLoci tool of the GATK (*see Methods*), and the lowest 2% average mapping quality in the global alignment. B) Non-overlapping regions remove from all within-host datasets: polymorphisms in the within-host dataset demonstrating strand-bias or tail-distance-bias in any of the 19 samples as identified by a previously described method [22] https://github.com/tamilieberman/IntrasamplePolymorphismCaller plus 5bp up- and down-stream.

**S13 File. H37Rv gene transfer format.** Tab-delimited gene transfer format file for H37Rv obtained from tbdb.org. Chromosome has been changed to reflect the reference sequence chromosome used for the between-host dataset.

**S14 Table. Polymorphisms passing filtering criteria for each within-host patient.** Each tab corresponds to a patient (patA-E) and is followed by either “subsampled” or “fst”. Tables list allele counts at polymorphic sites used in gene-wise (subsamples) or single nucleotide polymorphism (fst) analyses.

**S15 Functional Enrichment Categories.** We used the following annotation categories to classify M.tb genes: computationally predicted Clusters of Orthologous Groups (COG) (*n* = 21 categories) [40]; essential and nonessential genes for growth in vitro as determined by transposon site hybridization (TraSH) mutagenesis [107]; genes essential for growth in a murine model of TB [108]; "targets of independent mutation" associated with drug resistance [44]; and the M.tb-specific, manually curated functional annotation lists from TubercuList (*n* = 7) [39]. COG annotations for the H37Rv genome were obtained from the TB database (tbdb.org) [105], as were the TraSH “in vitro essential”, “in vivo essential”, and “nonessential” gene annotations. TubercuList functional annotations were obtained from tuberculist.epfl.ch and reflect the most up-to-date annotations when the database was accessed (12/01/2013). We did not include the TubercuList categories “PE/PPE”, “stable RNA”, and “unknown” in our analyses.

